# Neuroprogesterone signals through Src kinase within RP3V neurons to induce the luteinizing hormone surge in female rats

**DOI:** 10.1101/835470

**Authors:** Timbora Chuon, Micah Feri, Claire Carlson, Sharity Ondrejik, Paul Micevych, Kevin Sinchak

## Abstract

Neural circuits in female rats are exposed to estradiol and sequential progesterone to regulate the luteinizing hormone (LH) surge and thus ovulation. Estradiol induces progesterone receptors (PGRs) in rostral periventricular region of the third ventricle (RP3V) kisspeptin neurons, and positive feedback estradiol concentrations induce neuroprogesterone (neuroP) synthesis in hypothalamic astrocytes that signal to PGRs expressed in kisspeptin neurons to trigger the LH surge. We tested the hypothesis that neuroP-PGR signals through Src family kinase (Src) to trigger the LH surge. As in vitro, PGR and Src are co-expressed in RP3V neurons. Estradiol treatment increased the number of PGR immunopositive cells and PGR and Src colocalization. Infusion of the Src inhibitor (PP2) into the RP3V, attenuated the LH surge measured by ELISA in trunk blood collected 53 hours post-EB injection. While PP2 reduced the LH surge in 50 μg EB treated ovariectomized/adrenalectomized (ovx/adx) rats, activation of either PGR or Src in 2μg EB primed animals significantly elevated LH concentrations compared with DMSO treated ovx/adx rats. These results support the importance of Src in the estradiol and neuroP triggering of the LH surge.

## INTRODUCTION

Ovulation is coordinated with reproductive behavior and development of the uterine lining for copulation to result fertilization and pregnancy. In all mammals, and specifically for rodents, these events are mediated by hypothalamic-pituitary-ovarian axis hormones. Circulating estradiol concentrations slowly rise during diestrus I and II, priming luteinizing hormone (LH) surge neurocircuitry demonstrated by the increased expression of classical, “nuclear” progesterone receptor (PGR) and gonadotropin releasing hormone (GnRH) synthesis (Chappell & Levine, 2000). This initial rise of estradiol exerts a negative feedback on the hypothalamus and pituitary. As the rat cycle progresses into proestrus, estradiol concentrations rapidly rise inducing so-called estrogen positive feedback, in the afternoon, triggering the preovulatory LH surge that stimulating ovarian follicle ovulation and luteinization of the remaining follicular cells (Chazal, Faudon, Gogan, & Laplante, 1974). Positive feedback levels of estradiol induce hypothalamic astrocytes to synthesize neuroprogesterone (neuroP) that acts on induced PGR in neurons to release kisspeptin onto GnRH neurons triggering the LH surge (Mittelman-Smith, Wong, Kathiresan, & Micevych, 2015; Mittelman-Smith, Wong, & Micevych, 2018; Stephens et al., 2015; Zwain & Yen, 1999). Since the kisspeptin neurons that regulated the surge release of GnRH are located in the rostral periventricular region of the third ventricle (RP3V; (Clarkson & Herbison, 2006; Han et al., 2005; Liu, Lee, & Herbison, 2008; Wintermantel et al., 2006)), we hypothesized that neuroP acts on PGRs in these kisspeptin neurons to induce the LH surge (Delhousay, Chuon, Mittleman-Smith, Micevych, & Sinchak, 2019; Hu et al., 2015; Mittelman-Smith et al., 2018). This hypothesis is supported by experiments that demonstrated estradiol stimulated neuroP in hypothalamic astrocytes, without which there is no LH surge. Significantly, PGRs are required to initiate and reach the full magnitude and duration of the LH surge (Mahesh & Brann, 1998; Micevych, Matt, & Go, 1981), the LH surge cannot be induced in PGR knockout mice (Chappell et al., 1999; Mahesh & Brann, 1992; Micevych et al., 2003), and the removal of PGR from kisspeptin neurons causes infertility, the loss of puberty, and loss of the LH surge (Stephens et al., 2015).

Although PGR has long been classified as a transcription factor, it is one of a number of nuclear steroid receptors that can be trafficked to the plasma membrane and interact with other signaling proteins to initiate rapid signaling at the level of the plasma membrane (Boonyaratanakornkit et al., 2007; Boonyaratanakornkit et al., 2001; Boulware & Mermelstein, 2009). At the membrane, PGR complexes with (Src) (Boonyaratanakornkit et al., 2001). Src is a family member of nonreceptor tyrosine kinases (Pang, Wang, Valtorta, Benfenati, & Greengard, 1988), which interacts directly with PGR via its SH3 domain (Boonyaratanakornkit et al., 2001).

We have demonstrated PGR-Src interdependent interactions both *in vivo* and *in vitro.* The PGR-B isoform is complexed with Src in the plasma membrane within the arcuate nucleus of the hypothalamus (ARH), a region important for facilitation of lordosis (reviewed in Micevych & Sinchak, 2018a)). Inhibiting Src via ARH infusions of PP2 blocks progesterone facilitating of lordosis in estradiol-primed ovariectomized (ovx) rats, and blocking PGR prevents Src activator facilitation of lordosis (reviewed in (Micevych & Sinchak, 2018a)). An *in vitro* model of adult female hypothalamic kisspeptin neurons in the RP3V also supports PGR-Src interactions. Cultured mHypoA51 neurons express PGR and Src (Micevych, Wong, & Mittelman-Smith, 2015; Mittelman-Smith et al., 2018). Activation of Src in mHypoA51 neurons induced kisspeptin release while inhibiting Src activation blocks progesterone activation of MAPK and facilitated kisspeptin release, implying that progesterone and Src interact to stimulate kisspeptin (Mittelman-Smith et al., 2018). These results suggest that kisspeptin neurons utilize rapid, membrane-initiated signaling pathways induced by progesterone. Based on these results, the present study tested the hypothesis that neuroP acts through PGR that complexes with and signals through Src kinase to induce the release of kisspeptin in the RP3V, triggering the LH surge in ovx/adrenalectomized (adx) rats.

## MATERIALS AND METHODS

### Animals

In Experiment I, ovx rats were purchased from Charles River (Portage, MI) at 200-225 grams in weight or 55-65 days old. Ovx surgeries were performed by the supplier. Animals were housed in a light (12/12 light/dark cycle, lights on at 0600 hours) and climate-controlled room with food and water provided *ad libitum*. Wound clips were removed within one week of the arrival date.

In Experiments II-IV, ovx/adx rats were purchased from Charles River (Portage, MI) at 200-225 grams in weight or 55-65 days old. Ovx/adx surgeries were performed by the supplier. Drinking water was provided for the entirety of the experiment, consisting of NaCl (0.9%) and corticosterone (10 μg/ml; Sigma Aldrich; St. Louis, MO) (P. Micevych et al., 2003). Animals were housed in a light (12/12 light/dark cycle, lights on at 0600 hours) and climate-controlled room with food and water provided *ad libitum*. Rats were double-housed prior to cannulation. After cannulation surgeries, they were single-housed to prevent head-cap damage. Wound clips were removed within one week of the arrival date. All procedures involving animals were approved by the California State University, Long Beach IACUC.

Steroid/oil subcutaneous (s.c.) treatments were dissolved in safflower oil (oil) so that the volume of each injection was 0.1 ml.

### Experiment I. PGR-Src Double-Label Immunohistochemistry

To verify the colocalization of PGR and Src were within RP3V neurons (Figure 2), ovx Long Evans 200-225g rats were treated s.c. with either estradiol benzoate (EB; 2 μg) or oil once every four days for 3 cycles. Forty-eight hours after the third cycle, rats were deeply anesthetized with isoflurane and transcardially perfused with chilled 0.9% saline followed by PARA (4% paraformaldehyde in 0.1M Sorenson’s phosphate buffer) (Eckersell, Popper, & Micevych, 1998; Sinchak & Micevych, 2001). Brains were extracted, post-fixed in 4% paraformaldehyde for 24 hours, and cryoprotected. Brains were then sectioned in a cryostat at 20 μm thickness through the RP3V, and collected into phosphate-buffered saline (PBS, pH 7.5). Double-labeled immunohistochemistry was conducted using a polyclonal PGR primary antibody raised in rabbit (Table 1) that recognizes PGR-A and PGR-B, and a monoclonal primary antibody raised in mouse that recognizes Src 416-nonphosphorylated (Table 1). Free-floating brain sections were washed in PBS, followed by incubation in PBS containing 10% MeOH and 3% H2O2 for 10 minutes, then incubated in a nonspecific blocking solution containing PBS, 0.2% Triton X-100 (TX), 1% normal goat serum (NGS), and 1% bovine serum albumin (BSA) for 1 hour. Sections were incubated for an additional 48 hours at 4°C in the nonspecific blocking solution with a cocktail of PGR antibody (1:2000) and Src (1:800,000). Sections were washed in PBS and then tris-buffered saline (TBS, pH 7.5), and incubated for 2 hours in TBS containing 1% NGS, 0.2% TX and a biotinylated goat anti-rabbit secondary antibody conjugated with fluorescein (FITC) for PGR (1:200, Jackson ImmunoResearch Laboratories Inc., West Grove, PA), and a rhodamine-conjugated goat anti-mouse secondary antibody (TRITC) for Src (1:200, Jackson ImmunoResearch Laboratories Inc., West Grove, PA). Brain sections were rinsed in two final TBS washes, transferred to tris buffer, and mounted onto Superfrost Plus slides (Fisher Scientific). Mounted sections were dried on a slide warmer and cover-slipped using Aqua-Poly/Mount (Polysciences Inc., Warrington, PA). Visualization of immunohistochemistry was conducted via fluorescent microscopy (Leica DM6000, Leica Microsystems, Wetzlar, Germany) and Olympus Fluoview 1000 confocal laser scanning system (Olympus America Inc., Center Valley, PA).

**Table 1:** antibodies

| Antigen Target | Species raised in | Dilution | Company | Catalog No. |
| --- | --- | --- | --- | --- |
| PGR | Rabbit | 1:2,000 | Cell Signaling | D8Q2J |
| Src 416-NP | Mouse | 1:800,000 | Cell Signaling | 7G9 |

**FIGURE 1.**
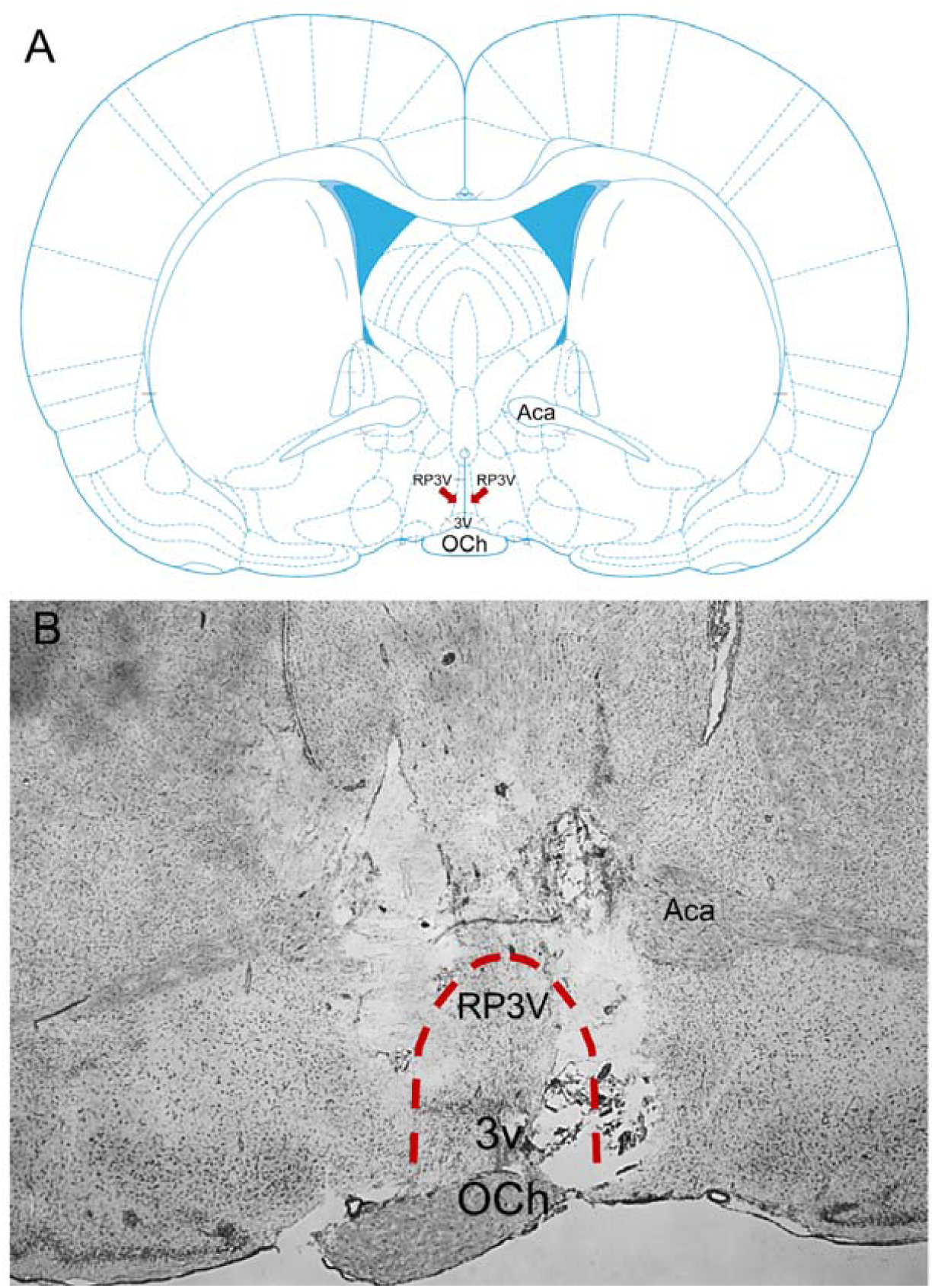
Region of site-specific infusions at the level of the rostral periventricular area of the third ventricle (RP3V; A) (Paxinos & Watson, 1998). Adult ovx/adx Long-Evans rats were implanted with a bilateral guide cannula aimed at the RP3V. Brains were coronally sectioned at 20μm and thionin stained to visualize cannula tract placement and infusion site (B). Och = optic chiasm, Aca = anterior commissure; 3v = third ventricle.

**FIGURE 2.**
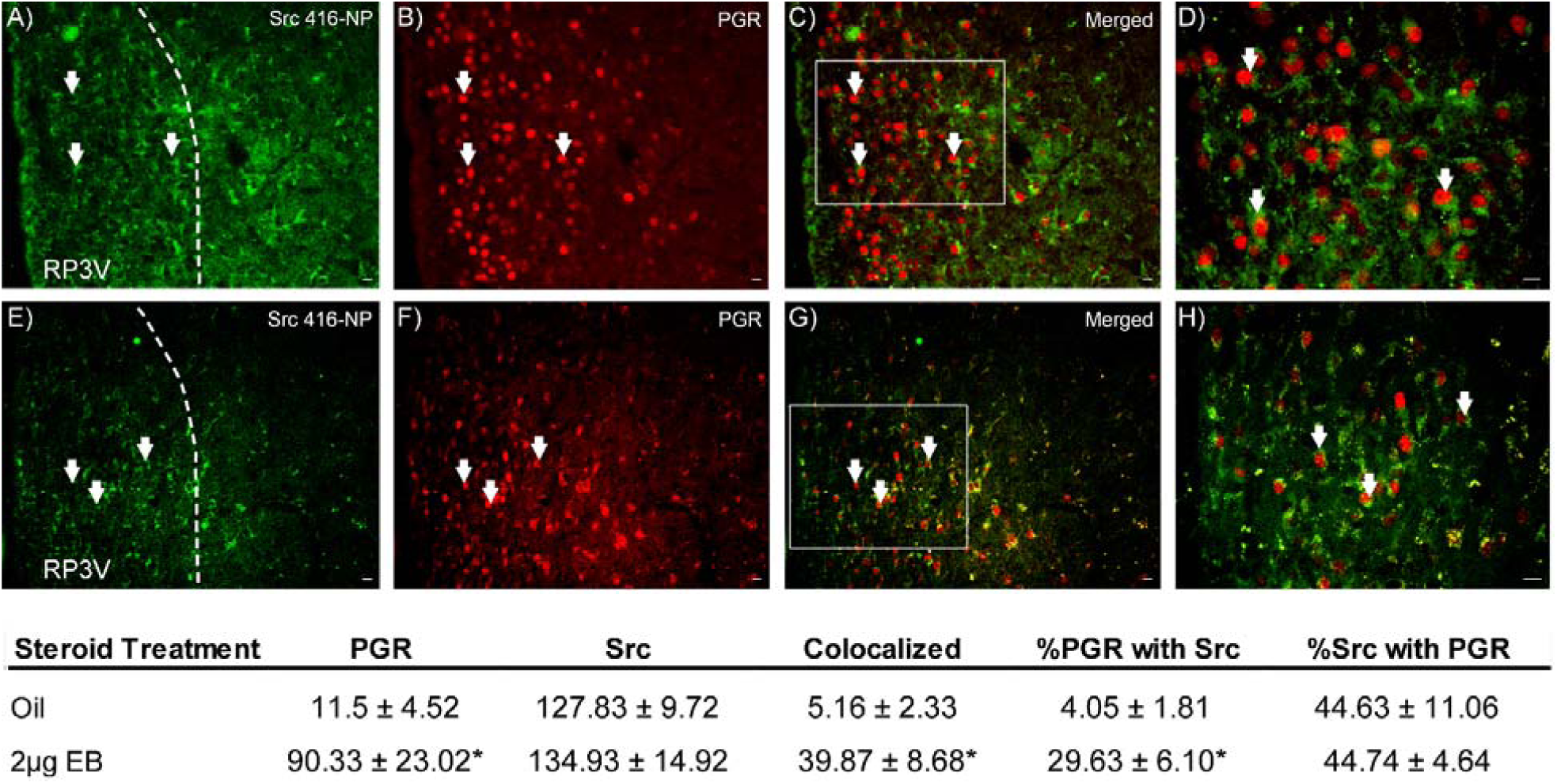
Estradiol increases PGR immunoreactivity and PGR-Src co-expression. Photomicrographs of immunohistochemical co-expression of classical progesterone receptor (PGR) and Src 416-nonphosphorylated (NP) in the RP3V of EB and oil-treated animals. PGR (Red, B and F) and Src (Green, A and E) immunopositive neurons are co-localized at the level of the RP3V (C and G, 40x photomicrographs D and H). Dotted outline indicates RP3V region where cells were counted. TRITC labeling of PGR immunoreactivity (arrows; red) was mainly localized in the nucleus of neurons. FITC labeling of Src was localized to the cytoplasm and extended to some neuronal processes (arrowheads; green). 3V = third ventricle. The number of immunopositive RP3V PGR neurons was significantly increased by EB treatment in comparison to oil, but the number of Src stained cells was not affected. The number of neurons that were immunopositive for both PGR and Src was increased by EB treatment in comparison to oil treatment. Further, the percentage of Src immunopositive cells that expressed PGR remain unchanged, but estradiol treatment significantly increases the percentage of PGR immunopositive cells that were stained positive for Src. * = significantly greater that Oil treatment group (p < 0.001).

### Immunohistochemistry Analysis

To confirm colocalization of immunoreactivity for PGR and Src, sections were analyzed by obtaining Z-stacks through the depth of the tissue within the RP3V. Three-dimensional reconstructions were used to determine colocalization of PGR and Src immunoreactivity (Mills, Sohn, & Micevych, 2004). FITC was visualized with a 465-495 nm emission filter and a 515-555 nm bandpass filter. TRITC was visualized with a 515-550 nm emission filter and 600-640 nm bandpass filter.

Analyses of single and double-labeled RP3V neurons were performed on photomicrographs of PGR and Src immunofluorescent staining in the RP3V using the ImageJ cell counter application (version 1.32j; National Institutes of Health, Bethesda, MD). Quantitative measurements of Src and PGR colocalization were performed by counting the number of RP3V neurons that expressed cytoplasmic staining for Src, nuclear staining for PGR, or both Src and PGR. To determine the proportion of Src or PGR immunoreactive neurons that co-express Src and PGR, a percentage was obtained by dividing the number of total Src and PGR colocalized neurons by either the total number of Src only or PGR only immunoreactive neurons, and then multiplying the calculated number by 100. All statistics for immunohistochemistry were performed using SigmaPlot 11.0. PGR and Src images of immunopositive RP3V neurons were taken with a DM6000 epiluminescent microscope. Colocalization was confirmed by using a Leica DFC 360FX monochrome digital camera and Leica AF-LAS microscope software to detect Src and PGR through the use of FITC and TRITC filter cubes, respectively.

### *Experiment II.* Duolink Proximity Ligation Assay

Results in Experiment I indicated that PGR and Src colocalization was increased with estradiol treatment (Figure 2). To determine PGR and Src interactions in the RP3V, and whether estradiol increases the interactions of PGR and Src, a Duolink Proximity Ligation Assay (PLA; Sigma Aldrich, Carlsbad, CA) analysis was performed on free-floating tissues from EB and oil-primed treated rats (described in Experiment I). For the Duolink PLA, tissue was incubated with primary antibodies that bind to PGR and Src and then secondary antibodies conjugated to oligonucleotides that to form a circular template if the distance between the oligonucleotides were less than 40 nm apart (Gullberg & Andersson, 2010). Complementary labeled, fluorescently tagged oligonucleotide probes were used to detect and amplify the PGR-Src proteins in close apposition for visualization by fluorescent confocal microscopy. Free-floating brain sections were washed in PBS prior to being incubated in a Duolink PLA kit blocking solution, and then incubated for 48 hours in PBS solution containing monoclonal Src antibody raised in mouse (Table 1) that detects nonphosphorylated Src at tyrosine-416 (1:800,000) and polyclonal PGR antibody raised in rabbit (1:2000; Table 1). Tissue sections were then processed according to the manufacturer’s instructions, transferred into tris buffer and then mounted onto Superfrost Plus slides (Fisher Scientific). Mounted sections were dried on a 37°C slide warmer and incubated in Hoechst (Sigma Aldrich). Mounted sections were dried and cover-slipped using Aqua-Poly/Mount (Polysciences, Inc., Warrington, PA). Visualization of Duolink PLA was conducted using fluorescent microscopy (Leica DM6000, Leica Microsystems, Wetzlar, Germany) and Olympus Fluoview 1000 confocal laser scanning system (Olympus America Inc., Center Valley, PA).

### Duolink PLA Analysis

With the Duolink PLA, extranuclear punctate red fluorescent staining represents a positive interaction of PGR and Src in close proximity. Analysis of positive PLA staining in the RP3V was performed using ImageJ (version 1.32j; National Institutes of Health, Bethesda, MD) cell counter to count the number of Hoechst stained nuclei with red, positive PGR-Src interactions in the RP3V. A total of three brain sections containing the RP3V per animal were counted to create an average number of PGR-Src interactions per animal (n = 4). All statistics for Duolink PLA were performed using SigmaPlot 11.0.

### Experimental Design – Experiments III-V Src activation of LH surge

#### Experiment III

To test that Src activation mediates EB-induction of the LH surge (Figure 4), adult Long Evans ovx/adx rats were implanted with bilateral cannulas (PlasticsOne; Roanoke, VA) aimed at the RP3V (Figure 3). At 21 days post ovx/adx surgery (P. Micevych et al., 2003) all animals were given EB (50μg/ 0.1 ml s.c.) at 1200 hours. On the same day (Day 0), all animals received bilateral RP3V infusions of either Src inhibitor (PP2, 50 nmol; Table 2) or DMSO and again on Days 1 and 2 at 1100 hours. On Day 2, animals were deeply anesthetized with isoflurane at 1730 hours (just prior to lights out in out colony), trunk blood was collected via decapitation, and brains were collected for confirming cannula placement. LH was measured in serum via enzyme-linked immunosorbent assay (ELISA) to determine whether Src inhibition blocked the estradiol-induced rise in LH concentrations (indicative of the LH surge). Results were analyzed by t-test with p ≤ 0.05 significance level.

**Table 2:** Drugs and steroids

| Steroid/Drug | Action | Vehicle | Concentration | Company | Catalog No. |
| --- | --- | --- | --- | --- | --- |
| Src Activator | Src family agonist | Saline | 50 nmol | Santa Cruz Biotechnology | SC-3052 |
| Progesterone | Progesterone receptor agonist | DMSO | 50 nmol | Sigma Aldrich | P0130 |
| R5020 | PGR-specific agonist | DMSO | 50 nmol | PerkinElmer | NLP004005MG |
| RU486 | Progesterone receptor inhibitor | DMSO | 50 nmol | Sigma Aldrich | M8046 |
| PP2 | Src family inhibitor | DMSO | 50 nmol | Tocris | 1407 |

**FIGURE 3.**
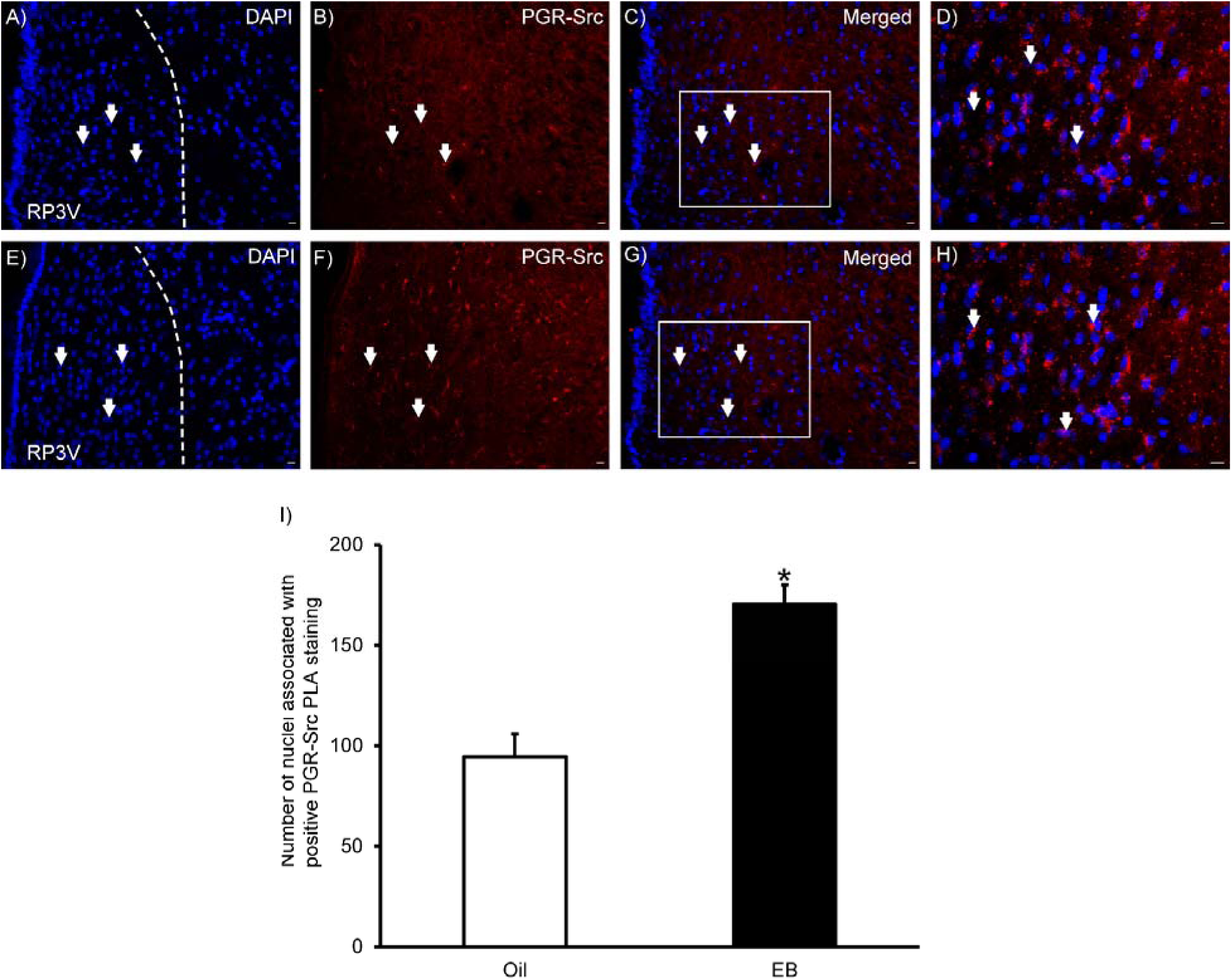
Estradiol increased the number of RP3V neurons with PGR and Src in close proximity. Photomicrographs of Duolink Proximity Ligation Assay (PLA) of classical progesterone receptor (PGR) and Src interactions in the RP3V of oil and EB-treated animals (3A-H). PGR and Src positive Duolink PLA interactions (red punctate staining) are within close apposition (< 40 nm) at the level of the RP3V. PGR-Src interactions surrounded Hoechst-stained nuclei (arrows). Estradiol treatment (3E-H) significantly increased the number of Hoechst-stained nuclei associated with positive PLA staining for PGR-Src (3I) in comparison to oil-treated animals (3A-D). * = significantly greater that Oil treatment group (p < 0.05).

**FIGURE 4:**
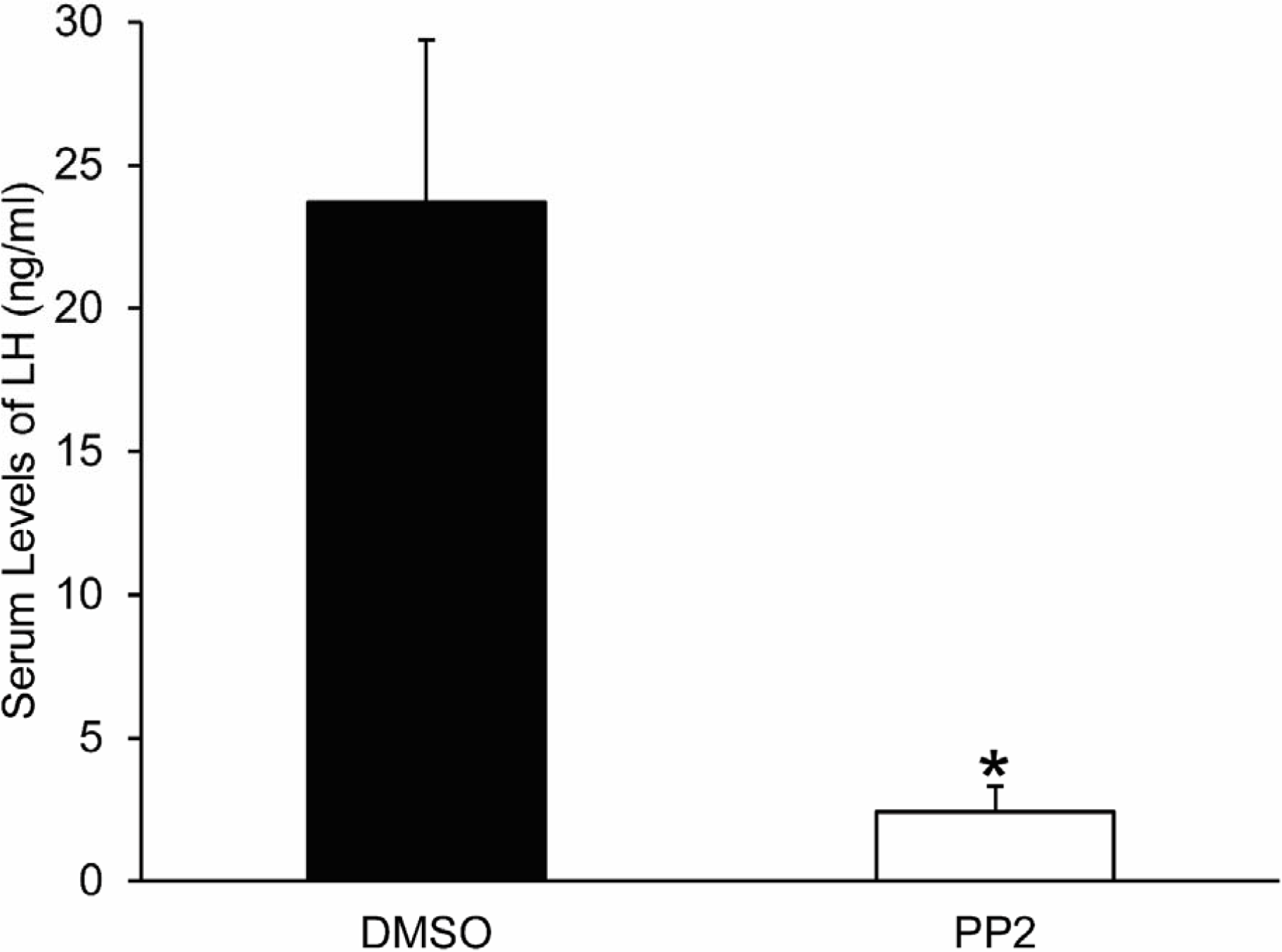
Inhibiting RP3V Src activation blocked the estradiol-induced LH surge in ovx/adx rats. All animals were treated with 50μg estradiol benzoate (EB) at 1200 hours and pretreated with either DMSO (n = 4) or Src antagonist (PP2) (n = 4) at 1100 hours, and on the next two days animals were infused daily with DMSO or PP2 and trunk blood was collected just prior to lights out in the colony on day 2. PP2 treatment significantly reduced serum LH concentrations compared to DMSO infused EB treated ovx rats. * = significantly greater than PP2 treatment (p < 0.05).

#### Experiment IV

Based on Experiment III results, we tested whether Src activation, like neuroP, triggers the LH surge in rats primed with a dose of subthreshold EB dose (s.c.). A 2 μg dose of EB upregulates PGR (Figure 5) (Chappell & Levine, 2000) but does not induce neuroP synthesis or an LH surge. Animal were treated as described in Exp III except they received EB (2μg/ 0.1 ml s.c.). Group 1 received DMSO at 1100 hours, followed by progesterone at 1530 hours. Group 2 received saline at 1100 hours, followed by Src activator (Table 2) at 1530 hours. Groups 3 and 4 received an initial infusion of either progesterone receptor antagonist (RU486; Table 2) or Src inhibitor (PP2; Table 2) at 1100 hours, followed by Src activator for both groups at 1530 hours. Group 5, the controls, received two infusions of DMSO at 1100 hours and 1530 hours. At 1730 hours, trunk blood was collected from all groups. Serum LH levels were measured by ELISA. Results were analyzed using a one-way ANOVA, followed by Student-Newman-Keuls (SNK) post hoc analysis.

**FIGURE 5.**
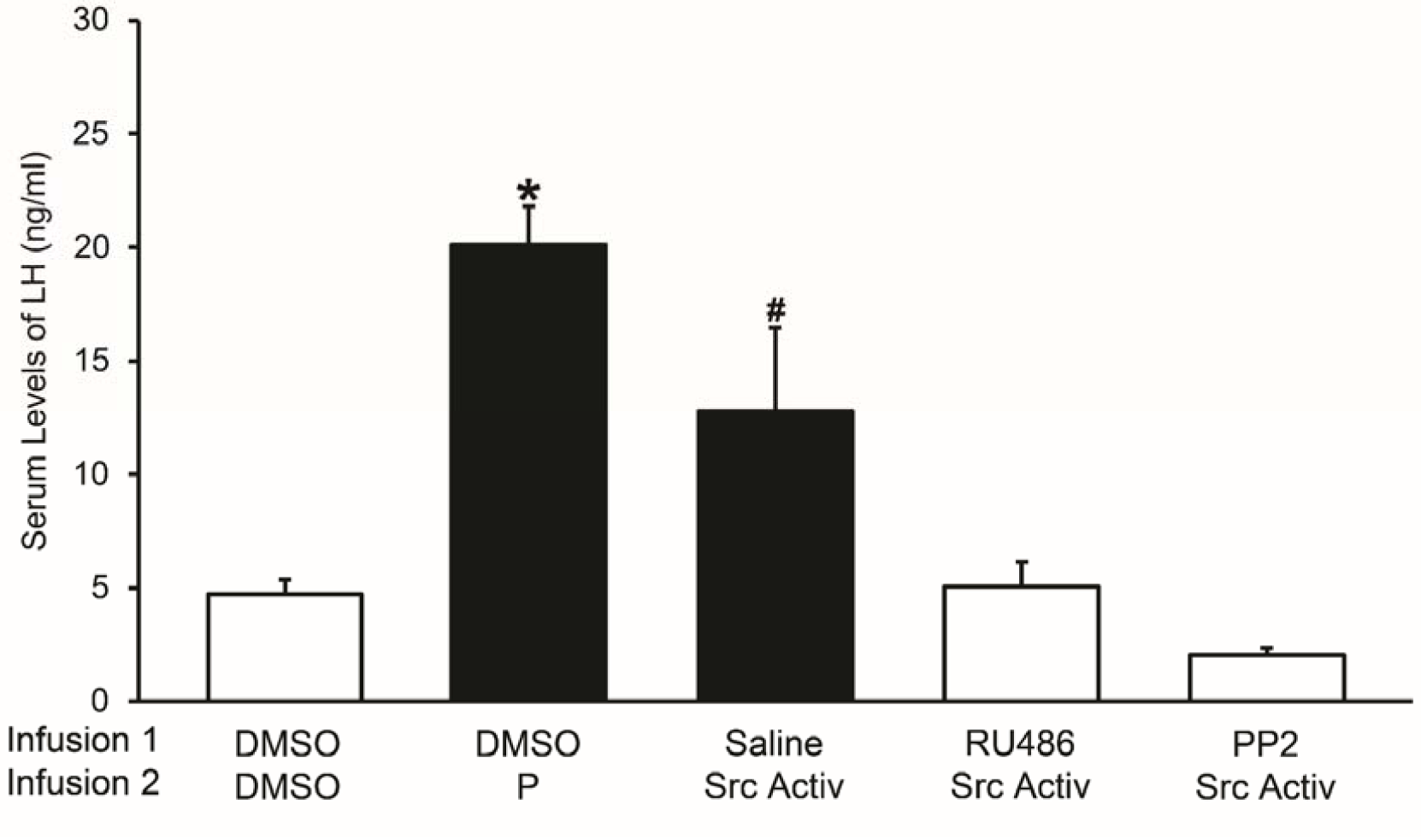
Interdependence of Src signaling with PGR to induce the LH surge. All animals were treated with 2μg EB and received two sequential infusions. The first set of RP3V infusions consisted of DMSO, Saline, RU486 or PP2 (n = 4 across all groups) administered 6.5 hours prior to blood collection. The second infusion administered was either DMSO, P, or Src Activator administered 2 hours prior to blood collection. DMSO + P and Saline + Src Activator had significantly higher mean LH concentrations compared to DMSO + DMSO, PP2 + Src Activator, and RU486 + Src Activator treated animals, and DMSO + P was significantly greater than Saline + Src Activator. * = significantly greater than all other treatment groups (p < 0.001). # = significantly different from all other treatment groups (p < 0.001).

#### Experiment V

To test whether Src activation is sufficient to induce the LH surge in 2μg EB-primed rats, adult ovx/adx Long Evans rats were implanted with a bilateral cannulae targeted at the RP3V. The first RP3V infusion was either DMSO, Src inhibitor (PP2), or PGR antagonist (RU486) at 1100 hours on Day 2 of the experiment. At 1530 hours, all animals received a sequential infusion of a PGR-specific agonist (R5020) and trunk blood was collected at 1730 hours. Serum LH levels were measured using ELISA, and results were analyzed using a one-way ANOVA followed by post hoc analysis by Holm-Sidak test for ANOVA p ≤ 0.05.

##### Stereotaxic Surgery for Implantation of Guide Cannulae

Animals were deeply anesthetized with isoflurane 13 days after ovx/adx surgery (2-3% in equal parts oxygen, Western Medical Supply Inc., Arcadia, CA), injected with an analgesic (Rimadyl 1 mg/ml; Western Medical Supply Inc., Arcadia, CA) and secured into a stereotaxic frame. Self-tapping bone screws were inserted into the skull to anchor the dental cement and cannula to the cranium. A bilateral cannula (22 gauge, 8 mm below pedestal; Plastics One, Roanoke, VA) was then implanted and aimed at the RP3V (coordinates from Bregma: 0° angle; anterior +0.1mm, lateral 0.7 mm, ventral −6.5mm, −3.3mm tooth bar, Figure 1). After dental cement secured the cannula to bone for attachment screws, a stylet was placed in the guide cannula, protruding no more than 0.5mm beyond the opening of the guide cannula to keep the cannula patent and covered with a head cap. After surgery, animals were single-housed and received oral antibiotics through the drinking water (0.5 mg/ml trimethoprim and sulfamethoxazole; TMS; Western Medical Supply Inc., Arcadia, CA). Animals were allowed to recover for 7 days prior to steroid treatment and drug infusion.

##### Steroid and Drug Treatments

Subcutaneous injections were performed using steroids (2μg EB/0.1ml, 50μg EB/0.1ml) dissolved in sterile safflower oil. Drugs were bilaterally infused into the RP3V. Src activator was dissolved in sterile saline (50 nmol) and PP2, PGR agonist (R5020), and progesterone were dissolved in sterile DMSO (50 nmol). The total volume infused per side was 0.5 μl. All infusions (vehicles and drugs) were performed with a sterile microinjector (Plastics One; Roanoke, VA) connected to a 25 μl Hamilton syringe. Tubing used for infusions were a thin plastic Tygon™ tubing at a rate of 1 μl per minute by a microliter syringe pump (Stoelting Co., Wood Dale, IL). Microinjectors did not protrude more than 2 mm beyond the opening of the guide cannulae and drug diffusion remained for 1 minute after infusion to allow the drug to diffuse away from the injection needle. Stylets were reinserted into the guide cannulae following microinfusion and animals were returned to their home cage until blood collection.

##### Serum Collection and Cannula Tract Confirmation

For Experiments III, IV and V, animals were deeply anesthetized with isoflurane and sacrificed by decapitation at 1730 hours on Day 22, 53 hours following EB treatment. Immediately following decapitation, trunk blood and brains were collected. Blood clotted at room temperature for 90 minutes and then centrifuged at 2000 × g for 15 minutes at room temperature. Serum was collected and stored at −80° C until analysis for LH by ELISA. Extracted brains were flash frozen on dry ice and stored at −80° C until sectioned for cannula placement. To visualize cannula placement, brains were mounted onto a chuck using HistoPrep medium, sectioned in the coronal plane at 20 μm through the RP3V, and directly mounted onto Superfrost™ Plus slides. Slides were allowed to dry on a slide warmer and stored at −80° C until thionin stained for visualization of cannula tract placement with brightfield microscopy (Figure 1).

##### Rat LH Sandwich ELISA

Serum LH concentrations were measured by rat LH sandwich ELISA (Shibayagi via BioVendor; Asheville, NC) (Delhousay et al., 2019). This assay was performed as specified in the Shibayagi via BioVendor kit protocol, with samples diluted to 5x concentration. Standards and samples were run in duplicate, and results of the ELISA were measured by a colorimetric microplate reader (VarioSkan 2.2; Thermo Scientific Inc., USA) at 450nm. For each sample, the mean was calculated, and the group means were statistically analyzed and graphed.

## RESULTS

### Experiment I

Although we expect some PGR immunoreactivity at the cell membrane, the predominate staining was associated with the nucleus (Figure 2A). Therefore, PGR nuclear-stained cells were used for estimating the number of PGR expressing neurons. Src positive staining was associated with both the cytoplasm and the plasma membrane (Figure 2B). Colocalization was visualized as red, nuclear PGR staining surrounded by green Src staining (Figure 2C). The number of Src immunoreactive RP3V neurons was greater than the number of PGR immunopositive neurons, independent of steroid treatment (Two-way ANOVA, F = 164.077, df = 1, 15, p < 0.001; Holm-Sidak EB t = 4.992, Oil t = 13.123, P < 0.001, Figure 2I). EB increased the number of RP3V PGR cells significantly compared to oil treatment (Two-way ANOVA, F = 58.537, df = 1, 15, P < 0.001; Holm-Sidak, t = 9.476, Figure 2I). In contrast, EB treatment did not alter the number of Src immunopositive cells compared with oil treatment (Holm-Sidak, t = 1.344; Figure 5A). However, EB treatment increased the number of PGR/Src expressing cells (t-test, P < 0.001, df = 6, t = 9.448; Figure 5B), and the percentage of Src cells that express PGR (t-test, p < 0.001, df = 6, t = 7.583; Figure 2K). The percentage of PGR immunopositive cells expressing Src (t-test, p = 0.969, df = 6, t = 0.0403, Figure 2L). This subpopulation of PGR/Src neurons with supports the hypothesis that neuroP may activate a RP3V PGR-Src signaling pathway to induce the LH surge.

### Experiment II

To verify the PGR-Src interactions, in the RP3V, Duolink PLA was performed to test whether PGR and Src are in close apposition (<40nm). Positive staining for PGR and Src in close apposition was observed as extranuclear punctate red fluorescent staining that was associated with a Hoechst counterstained nucleus (Figure 3H). A positive PGR-Src PLA staining was significantly greater in 2μg EB-treated rats compared to oil treated rats, indicating an increase in PGR-Src interactions (t-test, p < 0.001, df = 22, t = −18.066, Figure 3I).

### Experiment III

Results from Experiment I and II indicate that PGR and Src are co-expressed in RP3V neurons and interact. Thus, we tested whether Src activation is necessary to induce the LH surge. As expected, the 50μg EB treatment with DMSO infused-animals had serum concentrations of LH that were indicative of a surge (Delhousay et al., 2019; P. Micevych et al., 2003; P. Micevych, Soma, & Sinchak, 2008) (Figure 4). Src inhibition, with PP2 in RP3V, significantly decreased serum LH concentrations compared to DMSO-infused controls (t-test, df = 6; t = 3.511, p = 0.015). The attenuation of the LH surge via Src inhibition supports the hypothesis that Src mediates neuroP signaling that triggers the LH surge.

### Experiment IV addressed whether neuroP signaling required Src activation

RP3V progesterone treated EB-primed rats had surge concentrations of LH, while DMSO treated animals were significantly lower (one-way ANOVA, df = 4, 15, F = 20.221, p < 0.001; n = 4 animals per group; SNK post hoc test, p < 0.05, Figure 5). Src Activator also increased LH levels compared to DMSO (SNK p < 0.001). However, LH levels induced by progesterone were greater than those achieved with Src Activator (SNK p < 0.001). Pretreatment with either PGR antagonist (RU486) or Src inhibitor PP2) significantly reduced levels of LH induced by Src Activator (SNK p < 0.001). Together these results indicate a PGR-Src interdependent signaling mechanism underlying the LH surge.

### Experiment V tested whether

PGR activation was sufficient to induce Src signaling and the LH surge. Treatment with the PGR-selective agonist, R5020, produced significantly elevated levels of LH compared with DMSO (R5020, one-way ANOVA, df = 2, 9; F = 10.687, p < 0.05; Holm-Sidak p < 0.05, Figure 8). Pretreating with a Src inhibitor (PP2) or PGR antagonist (RU486) reduced R5020 induction of LH surge. These results demonstrate that a PGR-Src signaling pathway can trigger the LH surge.

## DISCUSSION

The major findings of these experiments are that 1) neuroprogesterone activates an interdependent RP3V PGR-Src signaling pathway that triggers the LH surge; 2) a subset of RP3V neurons co-express PGR and Src immunopositive staining; 3) PGR and Src are in close proximity to each other; 4) estradiol priming increased the number of RP3V neurons that co-express PGR and Src and increased the level of PGR and Src that were in close proximity. The estradiol-induced LH surge mediated by neuroprogesterone was attenuated by inhibiting RP3V Src activation (Figure 4). In congruence, RP3V Src activation in estradiol-primed rats produced serum surge levels of LH (Figure 5). Infusing the PGR-specific agonist (R5020) into the RP3V induced an LH surge in EB-primed rats, suggesting classical PGR mediate neuroP induction of the LH surge (Figure 6). PGR and Src signaling are interdependent since PGR antagonism with RU486 blocked Src activation, and Src inhibition blocked PGR activation by R5020 to induce the LH surge. Further, the PGR-B isoform of the receptor is likely mediating progesterone signaling since PGR-B interacts with the SH3 domain of Src and not PGR-A (Boonyaratanakornkit et al., 2007). Taken together, these data support the hypothesis that neuroP activates an RP3V PGR-Src signaling pathway to trigger the LH surge.

**FIGURE 6.**
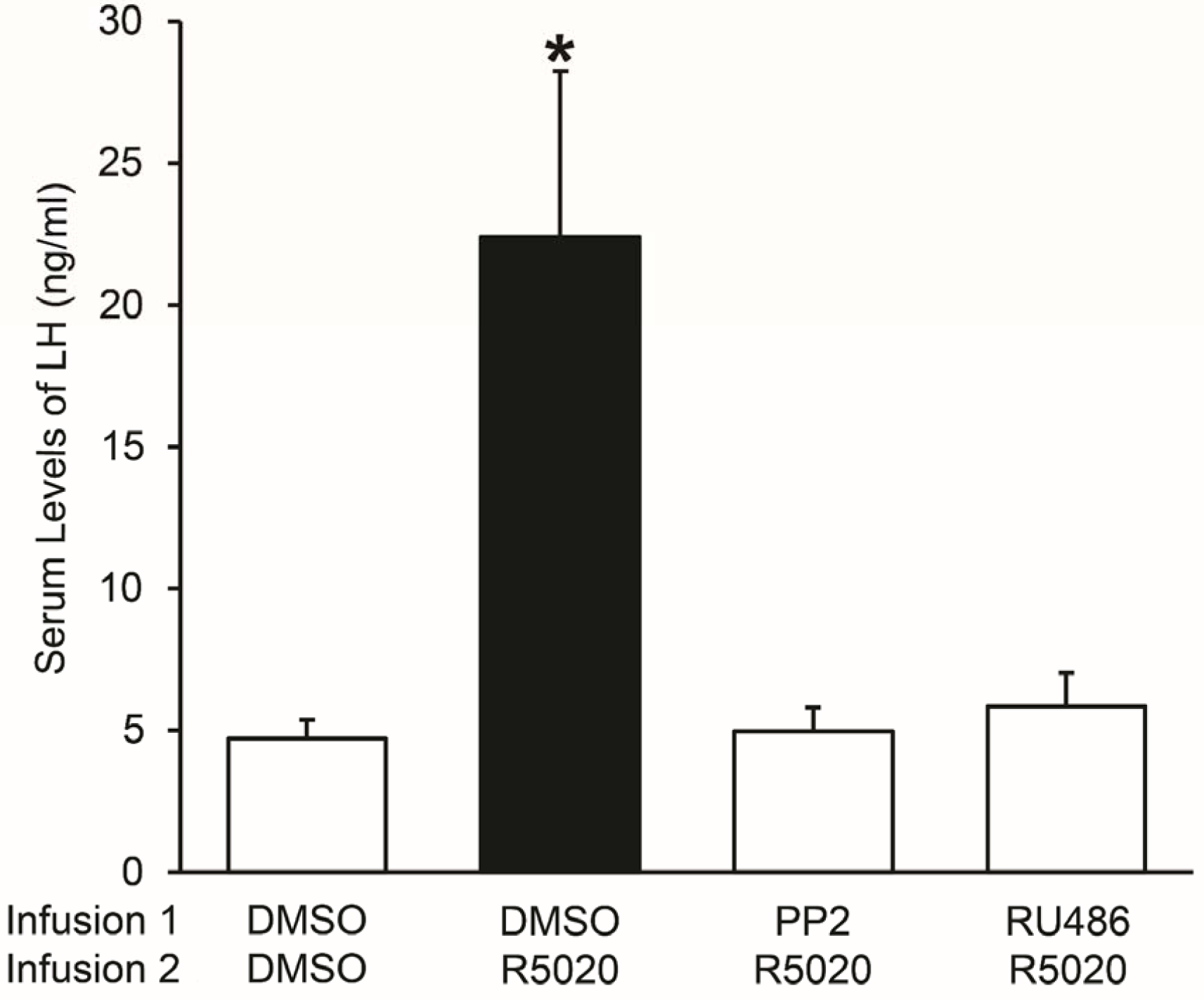
RP3V PGR mediate progesterone induction of the LH surge. RP3V infusion of PGR specific agonist, R5020, induces the LH surge in EB primed ovx/adx rats. All animals were treated with 2μg EB and received two sequential infusions. The first set of RP3V infusions (Infusion 1) was either DMSO, PP2, or RU486 (n = 4 across all groups) 6.5 hours prior to blood collection. The second infusion (Infusion 2) was either R5020 or DMSO administered 2 hours prior to blood collection. EB primed rats that received RP3V infusions of DMSO + R5020 had a significantly higher mean serum LH concentration compared to animals that received PP2 or RU486 treatment prior to R5020. * = significantly greater than Oil group (p < 0.05).

Neuroprogesterone induction of the LH surge is mediated by the release of kisspeptin from RP3V neurons (Delhousay et al., 2019) that requires the estradiol priming and the actions of neuroP (Mittelman-Smith et al., 2015; Mittelman-Smith et al., 2018). Our present *in vivo* findings indicate that neuroP activates a PGR-Src pathway in the RP3V region to induce the LH surge, presumably at the level of kisspeptin neuron. These conclusions are supported by previous findings in cultured kisspeptin neurons (mHypoA51) derived from adult female mice that are a model for RP3V kisspeptin neurons (Mittelman-Smith et al., 2015; Mittelman-Smith et al., 2018). The mHypoA51 neurons express both PGR and Src, and estradiol-priming increases PGR expression as well as increasing PGR-A and –B levels at the plasma membrane (Mittelman-Smith et al., 2015). Further, in co-culture experiments, estradiol induced neuroprogesterone from hypothalamic astrocytes induced kisspeptin release from mHypoA51 neurons (Mittelman-Smith et al., 2018). Src activation produced a similar release of kisspeptin from mHypoA51 neurons, supporting the idea that neuroprogesterone is signaling rapidly through a PGR-Src complex located in the plasma membrane that activates a MAPK pathway (Mittelman-Smith et al., 2018). Further support for neuroP signaling through plasma membrane PGR-Src complexes to trigger the LH surge is that estradiol increased the co-expression and extranuclear close proximity of PGR and Src in RP3V neurons. RP3V kisspeptin neurons *in vivo* express PGR (Clarkson, d’Anglemont de Tassigny, Moreno, Colledge, & Herbison, 2008), and we demonstrated that PGR and Src were colocalized in RP3V neurons and the number of colocalized cells was increased by estradiol priming. These data further suggest that a subpopulation of RP3V kisspeptin neurons to express both PGR and Src.

Our double-label immunohistochemistry revealed a subpopulation of RP3V neurons that co-express PGR and Src (Figure 2). As expected, estradiol priming increased the number of RP3V neurons that had positive nuclear staining for PGR compared to oil controls (Figure 2I). Although the increase in PGR neurons was determined by counts of positive nuclear PGR staining, it is likely that estradiol-induced upregulation of PGR on the membrane as well. In mHypoA51 neurons, estradiol treatment increased levels of PGR in a membrane biotinylation study (Mittelman-Smith et al., 2018). Further, preliminary data from our laboratory using cell fractionation and western blot technique reveals that 30 hours after estradiol priming, PGR levels in the plasma membrane increase (unpublished data). Thus, in vivo, it is likely that estradiol priming increases PGR levels in the plasma membrane of RP3V neurons. In contrast, the number of immunopositive Src (416-NP) in RP3V neurons was not increased by estradiol priming (Figure 2I), and the number of Src neurons appear to be greater than PGR neurons. Nonetheless, estradiol priming also significantly increased the number of RP3V neurons that were positively stained for both PGR and Src (Figure 2J). This suggests that estradiol priming increases levels of PGR-Src for subsequent neuroP/progesterone signaling to induce the LH surge. Interestingly, estradiol did not alter the percentage of PGR neurons that express Src (Figure 2K). However, the percentage of Src neurons that expressed PGR was significantly increased (Figure 2L). The increased percentage of Src neurons that expressed PGR was due to the estradiol induction of PGR in Src neurons since the number of Src neurons remain unaffected by estradiol treatment. These results support the idea that PGR-Src signaling is possible in a subset of RP3V neurons, and estradiol upregulates the PGR-Src signaling pathway. Since 40-60% of RP3V kisspeptin neurons co-expressed with PGR (Clarkson et al., 2008; Smith, Popa, Clifton, Hoffman, & Steiner, 2006), this supports the notion that neuroP acts directly on kisspeptin neurons via PGR-Src signaling to trigger the LH surge.

Our data also indicate that PGR and Src form signaling complexes, and estradiol increases these complexes in the RP3V. Duolink PLA demonstrates that a subpopulation of RP3V neurons have PGR and Src in close proximity, and that estradiol increased the levels of positive staining for PGR and Src in close proximity (Figure 3G) compared to oil-treated animals (Figure 3C). These results suggest that not only are PGRs upregulated in response to estradiol, but estradiol also increases PGR interactions with Src.

In previous studies, using a lower, 2μg priming dose of estradiol upregulates PGRs but is not sufficient to stimulate neuroP synthesis or the LH surge (Zwain & Yen, 1999). In an estradiol primed rat, infusion of progesterone or kisspeptin induced an LH surge (Delhousay et al., 2019). In this study we demonstrated that infusion of Src activator into the RP3V was sufficient to induce the LH surge in similarly estradiol-primed rats (Figure 5). Although Src activation significantly elevated LH concentrations indicative of an LH surge, LH concentrations were significantly lower than progesterone RP3V infusions that inducted the LH surge (Figure 5). The increase in LH concentration induced by RP3V Src activation was blocked by pretreating with either Src inhibitor (PP2) or PGR antagonist (RU486) (Figure 5). Similarly, PGR activation in RP3V neurons by R5020 increased LH concentrations that was blocked by pretreating with either PP2 or RU486 (Figure 6). These results suggest that neuroP is indeed signaling through Src in kisspeptin neurons to induce kisspeptin release and thus trigger the LH surge. The ability of RU486 to block Src induction of the LH surge, and the ability of PP2 to block PGR induction of the LH surge suggests that PGR and Src signaling is interdependent and not in serial or parallel. Although Src signaling is necessary for PGR induction of the LH surge, since RP3V infusion of progesterone induced greater LH levels than Src activator, it is possible that progesterone activates another pathway in parallel that is not sufficient to trigger the LH surge on its own, but may augment neuroP induction of the LH surge. It is possible that a portion of neuroP signaling that triggers the LH surge mediated by a family membrane progesterone receptors that are also expressed in kisspeptin neurons in vivo and in vitro (Mittelman-Smith et al., 2018; Thomas & Pang, 2012).

Our results demonstrating the interdependence of PGR and Src signaling that mediate neuroP induction of the LH surge are similar to *in vitro* mHypoa51 studies and ARH behavior experiments, suggesting that neuroP is signaling through a PGR-Src complex in the RP3V to trigger the LH surge (reviewed in Micevych & Sinchak, 2018b). Previous findings in our laboratory that demonstrate progesterone activates PGR-Src signaling in the ARH that rapidly facilitates sexual receptivity (Micevych & Sinchak, 2018a). These studies have demonstrated rapid, interdependent signaling of PGR and Src in the ARH that facilitate lordosis. Sexual receptivity is inhibited when Src activator was administered in the presence of a PGR antagonist, RU486 (reviewed in Micevych & Sinchak, 2018a). Conversely, inhibition of Src within the ARH blocked progesterone facilitation of lordosis. The similar pattern of interdependence of PGR and Src signaling to induce the LH surge that was observed in Experiment III. In the ARH, our laboratory has demonstrated that through co-immunoprecipitation experiments, the potential for PGR-Src complex formation localized to both the plasma membrane and the cytoplasm within the ARH (reviewed in Micevych & Sinchak, 2018a). Double labeled immunohistochemistry demonstrated that RP3V neurons co-express PGR and Src. Using a Duolink PLA demonstrated that PGR and Src are extranuclear and in close proximity to each other within the RP3V (Figure 3). Further, estradiol priming appears to upregulates the levels of PGR and Src that in close proximity and therefore increases the potential for formation of PGR-Src signaling complexes (Figure 3). Taken together, these double label immunohistochemical and Duolink assays further support the possibility that neuroP is signaling through a similar ARH interdependent PGR-Src complex in RP3V kisspeptin neurons to trigger the LH surge.

Although these results do not demonstrate that the PGR-Src signaling is localized to kisspeptin neurons, other findings indicate that this is the case. PGRs have been shown to be expressed in RP3V kisspeptin neurons (Clarkson, d’Anglemont de Tassigny, Colledge, Caraty, & Herbison, 2009). Mice that have PGRs knocked out specifically in kisspeptin neurons (kissPRKOs) lack an LH surge, demonstrating that PGR expression in kisspeptin neurons is essential for induction of the LH surge (Stephens et al., 2015). RP3V kisspeptin signaling is downstream of neuroP signaling (Delhousay et al., 2019). *In vitro* mHypoA51 neurons, a model of RP3V kisspeptin neurons, express Src and membrane-associated PGRs (Mittelman-Smith et al., 2018). Further, stimulation with either progesterone or PGR-specific agonist (R5020) induced rapid Src phosphorylation, and activation of Src stimulates the release of MAPK and kisspeptin (Mittelman-Smith et al., 2018). Therefore, PGR-Src signaling is likely to be acting directly on RP3V kisspeptin neurons that mediate the neuroP triggering of the LH surge. Further, these data support the idea that PGR is signaling through Src in a similar signaling mechanism as PGR-Src ARH neurocircuitry.

In summary, in our model for neuroP induction of the LH surge, initial estradiol priming effects increase PGR expression in kisspeptin neurons, while higher doses of estradiol upregulate neuroP synthesis secreted from hypothalamic astrocytes (Figure 7). Estradiol further upregulates PGR-Src colocalization and interactions in close proximity within the RP3V. During positive feedback, peaking levels of estradiol on proestrus induces the synthesis neuroP, which then signals through PGR-Src pathway. The PGR-Src complexes are likely to be in RP3V kisspeptin neurons to stimulate kisspeptin neurotransmission that triggers the LH surge. Together, these experiments support the hypothesis that Src and PGR interdependent signaling in the RP3V is necessary to trigger the LH surge. Although Src activation is sufficient to trigger the LH surge, progesterone may be activating other pathways in parallel to augment the surge since progesterone produced a greater magnitude of LH concentrations. These findings support the idea that neuroP signals through a PGR-Src complex, as observed in the ARH, to stimulate kisspeptin neurotransmission triggering the LH surge and inducing ovulation and luteinization of follicular cells.

**FIGURE 7.**
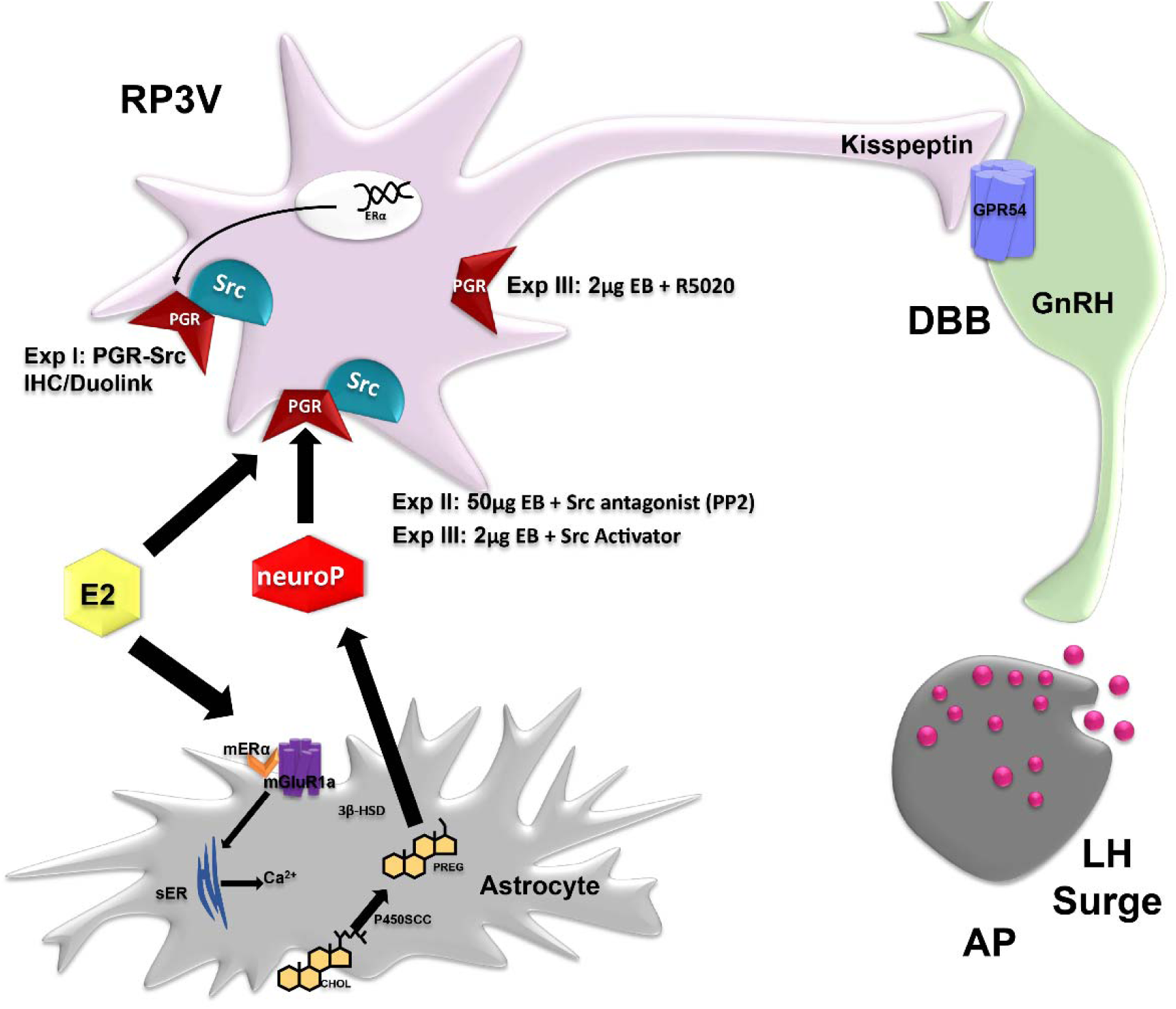
Proposed circuit of estradiol-induced neuroP on hypothalamic cells. In hypothalamic astrocytes, proestrous levels of estradiol (E2) act on membrane estrogen receptor-α (mERα) to increase intracellular calcium concentrations (Ca^2+^) by complexing with and signaling through the metabotropic glutamate receptor-1a type (mGluR1a). This releases Ca^2+^from the smooth endoplasmic reticulum (sER). In the mitochondrion, P450scc converts cholesterol (CHOL) to pregnenolone (PREG), which is further converted to neuroP by 3β-HSD. The newly synthesized neuroP activates E2-induced progesterone receptors (PGR) complexed with Src family kinase (Src) in AVPV kisspeptin neurons that project to and stimulate diagonal band of Broca (DBB) GnRH neurons, triggering the LH surge release from the anterior p itu itary (AP). Experiment I demonstrated inhibition of Src attenuated the LH surge in the presence of neuroP. Experiment II showed Src activation in the absence of neuroP can induce the LH surge, suggesting Src is downstream of neuroP signaling. Experiment III demonstrated PGR-Src colocalization and membrane localization and close proximity via immunohistochemistry (IHC) and Duolink, respectively. Modified from (Micevych et al., 2015).

## Acknowledgements

Authors thank Reema Tominna, Nasir Khan, Joshua Razon, Stephanie Huerta, Monica Eskander, Maxwell LaForest, Jessica Phan, Sima Chokr, and Julia Rodman for their technical assistance. We thank Dr. Seema Tiwari-Woodruff for use of her confocal microscope and Hana Yamate-Morgan for her technical assistance at the University of California, Riverside. Research was supported by NIH grant HD042635 (PM) HD058638 (KS) 5R25GM071638 (NIH RISE – CSULB), NIH grants HD04612 and the Doris A. Howell foundation.

